# Microstructural Properties of the Cerebellar Peduncles in Children with Developmental Language Disorder

**DOI:** 10.1101/2023.07.13.548858

**Authors:** Salomi S. Asaridou, Gabriel J. Cler, Anna Wiedemann, Saloni Krishnan, Harriet J. Smith, Hanna E. Willis, Máiréad P. Healy, Kate E. Watkins

**Affiliations:** Department of Experimental Psychology, Wellcome Centre for Integrative Neuroimaging, University of Oxford, Oxford, UK; Department of Speech & Hearing Sciences, University of Washington, Seattle, USA; Department of Psychiatry, University of Cambridge, Cambridge, UK; Department of Psychology, Royal Holloway, University of London, Egham Hill, Surrey, UK; MRC Cognition and Brain Sciences Unit, University of Cambridge, Cambridge, UK; Nuffield Department of Clinical Neuroscience, University of Oxford, Oxford, UK; Department of Psychology, University of Cambridge, Cambridge, UK

**Keywords:** Developmental language disorder, cerebellum, tractography, DTI, NODDI

## Abstract

Children with developmental language disorder (DLD) struggle to learn their native language for no apparent reason. While research on the neurobiological underpinnings of the disorder has focused on the role of cortico-striatal systems, little is known about the role of the cerebellum in DLD. Cortico-cerebellar circuits might be involved in the disorder as they contribute to complex sensorimotor skill learning, including the acquisition of spoken language. Here, we used diffusion-weighted imaging data from 77 typically developing and 54 children with DLD and performed probabilistic tractography to identify the cerebellum’s white matter tracts: the inferior, middle, and superior cerebellar peduncles. Children with DLD showed lower fractional anisotropy (FA) in the inferior cerebellar peduncles (ICP), fiber tracts that carry motor and sensory input via the inferior olive to the cerebellum. Lower FA in DLD was driven by lower axial diffusivity. Probing this further with more sophisticated modeling of diffusion data, we found higher orientation dispersion but no difference in neurite density in the ICP of DLD. Reduced FA is therefore unlikely to be reflecting microstructural differences in myelination in this tract, rather the organization of axons in these pathways is disrupted. ICP microstructure was not associated with language or motor coordination performance in our sample. We also found no differences in the middle and superior peduncles, the main pathways connecting the cerebellum with the cortex. To conclude, it is not cortico-cerebellar but atypical olivocerebellar white matter connections that characterize DLD and suggest the involvement of the olivocerebellar system in speech acquisition and development.

## 1. Introduction

Children with Developmental Language Disorder (DLD) show significant and unexplained deficits in producing or comprehending language or both compared with their peers (Bishop et al., 2016). A child with DLD can present with a wide range of problems including problems in speech sound discrimination and phonology, in word learning and vocabulary, expressive and receptive grammar (Conti-Ramsden et al., 2001; Krishnan et al., 2021), as well as pragmatics (Bishop, 2014) (see Bishop et al., 2016, 2017 for detailed description and terminology). Despite it being a highly prevalent disorder, research on the neurobiological basis of DLD is still scarce. The majority of neuroimaging studies conducted so far have highlighted differences between DLD and typically developing children (TD) in the function and structure of cortical structures (Mayes et al., 2015), in particular inferior frontal and superior temporal areas that are considered key nodes of the language (Fedorenko & Thompson-Schill, 2014). Among the most consistent findings in the literature is under-activation in left frontal and temporal cortical areas while processing language as well as microstructural differences in dorsal anatomical pathways that connect them (for a review see Asaridou & Watkins, 2022).

Another relatively consistent finding has been atypical structure in the basal ganglia (Badcock et al., 2012; Lee et al., 2013; Watkins et al., 2002). The basal ganglia play an important role in procedural learning, including speech motor learning during language acquisition (Karuza et al., 2013) and the ability to extract sequential regularities required for learning phonology and grammar (Krishnan et al., 2016). It has been hypothesized that the problems in DLD stem from deficient domain-general implicit learning mechanisms that involve the basal ganglia (Krishnan et al., 2016; Ullman et al., 2020). Children with DLD show poorer performance in non-linguistic procedural learning, particularly in sequence-based tasks and probabilistic categorical learning (Gabriel et al., 2013; Hsu & Bishop, 2014; Lee & Tomblin, 2012). Supporting this hypothesis, a recent study using quantitative MRI revealed differences in striatal myelin in children with DLD (Krishnan et al., 2022).

Cortico-cerebellar circuits, alongside cortico-striatal systems, contribute to complex sensorimotor skill learning, including the acquisition of spoken language (Ziegler & Ackermann, 2017). The cerebellum, together with frontal and temporal cortical areas and the basal ganglia, is part of a *FOXP2*-dependent circuitry that has been proposed to support speech motor control (Vargha-Khadem et al., 2005). Neural expression of *FOXP2*, a gene in which a point mutation results in a speech and language disorder in members of the KE family, has been found in both the basal ganglia and the cerebellum (Lai et al., 2003). Affected members of the KE family show atypical basal ganglia structure (Vargha-Khadem et al., 1998; Watkins et al., 2002) as well as atypical cerebellar structure (Argyropoulos et al., 2019). Children with cerebellar agenesis or congenital cerebellar malformations show delayed speech acquisition (Glickstein, 1994; Steinlin, 1998). Cerebellar mutism (lack of speech) and dysarthria can follow brain surgery that directly or indirectly affects the cerebellum in children (Küper & Timmann, 2013). Disruption of cerebro-cerebellar circuits, particularly during development, has also been documented in developmental disorders such as autism and dyslexia (for a review see Stoodley, 2016).

The role of the cerebellum in DLD has received little attention. This is partly because children with DLD perform as well as TD in tasks such as eyeblink conditioning, which rely on the cerebellum (Hardiman et al., 2013). Performance in other tasks, however, that involve the cerebellum (Miall & Christensen, 2004) and require fine motor skills does differ, with children with DLD performing worse than TD (Brookman et al., 2013; Jäncke et al., 2007; Powell & Bishop, 2008; Zelaznik & Goffman, 2010).

Given the uniform architecture of the cerebellum, functional specialization for language must be determined by inputs and outputs from peripheral sensors and effectors along with cortical and subcortical language circuits (Skipper & Lametti, 2021; Steele & Chakravarty, 2018). These inputs and outputs are mediated by the three main white matter pathways that connect the cerebellum to the rest of the brain known as the peduncles. The inferior and middle cerebellar peduncles provide cerebellar inputs, and the superior cerebellar peduncles deliver cerebellar outputs. Click or tap here to enter text.The inferior cerebellar peduncles consist of climbing fibers carrying predominantly afferent input from the inferior olive to the cerebellum (Moberget & Ivry, 2016). The middle cerebellar peduncles consist of mossy fibers carrying afferent input from the cortex to the cerebellum via the pons (Stoodley & Schmahmann, 2010). Lastly, the superior cerebellar peduncles consist of mossy fibers carrying efferent input from the cerebellum to the cortex via the thalamus (Moberget & Ivry, 2016). In typically developing children, the development of the middle and superior cerebellar peduncles peaks in early adolescence (12-15 years) while the inferior cerebellar peduncle can peak later, in mid adolescence (12-17 years) (Simmonds et al., 2014).

The role of the cerebellar white matter connections in speech and language remains relatively understudied. White matter in the cerebellar peduncles has been associated with reading skills in children and adolescents (Bruckert et al., 2020; Travis et al., 2015), and with speech rate in adults who stutter (Jossinger et al., 2021), while differences in white matter microstructure of the peduncles have been reported between controls and adults and children who stutter (Connally et al., 2014; Johnson et al., 2022). There is some evidence that cortico-cerebellar white matter connections show different microstructural properties in adolescents (>14 yrs) and young adults with DLD compared with a control group (Lee et al., 2020). Very little is known about cerebellar structure in children with DLD and, to the best of our knowledge, cerebellar connectivity has not yet been tested in this population.

The aim of the current study was to investigate cerebellar white matter connectivity in a large sample of children with DLD. We hypothesized that microstructural properties of the main cerebellar pathways (the inferior, middle, and superior cerebellar peduncles) will differ between children with DLD and TD children. We also hypothesized that microstructural characteristics of the cerebellar peduncles in our sample will be associated with performance on language and fine-motor tasks.

## 2. Methods

### 2.1 Participants

One-hundred-and-seventy-five children aged 10;0 – 15;11 (years; months) were recruited as part of the Oxford Brain Organisation and Language Development (OxBOLD) study. The study protocol was approved by the University of Oxford’s Medical Sciences Interdivisional Research Ethics Committee in accordance with the Declaration of Helsinki. Informed written consent was obtained from parents/guardians as well as from the child prior to study enrolment. Five participants did not complete the behavioral testing or imaging, seven were subsequently found not to meet the study’s inclusion criteria, three were excluded due to incidental findings, and one participant was excluded because their cerebellum was not included in the scans (not part of Field-Of-View). Complete data were therefore available for 159 participants with sample characteristics shown in Table 1.

**Table (1):** Sample characteristics of the OxBOLD study.

|  | TD (N=77) | DLD (N=54) | HSL (N=28) | *p* value |
| --- | --- | --- | --- | --- |
| **Demographics** |  |  |  |  |
| Age (in years) |  |  |  | 0.894 |
|  | 12.53 (1.72) | 12.40 (1.85) | 12.39 (1.56) |  |
| Gender |  |  |  | 0.032 |
| Male | 44 (57.1%) | 39 (72.2%) | 23 (82.1%) |  |
| Female | 33 (42.9%) | 15 (27.8%) | 5 (17.9%) |  |
| Handedness |  |  |  | 0.924 |
| Right | 66 (85.7%) | 45 (83.3%) | 24 (85.7%) |  |
| Left | 11 (14.3%) | 9 (16.7%) | 4 (14.3%) |  |
| **Neuropsychological Assessments** |  |  |  |  |
| Non-verbal IQ (mean = 10, SD = 3) |  |  |  | < 0.001 |
|  | 12.30 (1.90) | 8.66 (2.65) | 11.32 (1.94) |  |
| Language factor (a.u.) |  |  |  | < 0.001 |
|  | 0.77 (0.46) | -1.06 (0.60) | 0.00 (0.50) |  |
| Memory factor (a.u.) |  |  |  | < 0.001 |
|  | 0.54 (0.80) | -0.75 (0.75) | 0.05 (0.73) |  |
| Pegboard (z-score) |  |  |  | < 0.001 |
|  | -0.41 (0.85) | -1.56 (1.22) | -0.87 (0.85) |  |
| **Diffusion MRI Measures** |  |  |  |  |
| Whole brain FA |  |  |  | 0.052 |
|  | 0.24 (0.01) | 0.24 (0.01) | 0.24 (0.01) |  |
| SNR |  |  |  | 0.229 |
|  | 39.12 (5.02) | 37.54 (5.32) | 38.21 (5.57) |  |
| Relative Motion Displacement (mm) |  |  |  | 0.060 |
|  | 0.30 (0.12) | 0.33 (0.16) | 0.26 (0.09) |  |

*Abbreviations:* TD = typically developing, DLD = developmental language disorder, HSL = history of speech-language problems, FA = fractional anisotropy, SNR = Signal-to-Noise Ratio, a.u. = arbitrary units for factor scores. Note: *p*-values reported on group differences with One-Way ANOVA and Chi-square tests.

In order to be included in the study, participants had to pass a bilateral pure tone audiometric screening test to confirm normal hearing and score no more than two standard deviations below the normative mean in two non-verbal IQ tests, the WISC-IV Matrix Reasoning and Block Design Test (Wechsler, 2003). Participants were excluded if they had a diagnosis of a developmental disorder such as Down or William Syndrome, or a history of neurological impairments or neurological disorders such as epilepsy. Participants with a diagnosis of autistic spectrum disorder (ASD) or attention deficit hyperactivity disorder (ADHD) were also excluded. Similarly, children who scored more than seven on the hyperactivity subscale of the Strengths and Difficulties Questionnaire (SDQ; Goodman, 1997), or more than 15 on the Social Communication Questionnaire – Lifetime (SCQ; Rutter et al., 2003) were also excluded from the study. All participants grew up in the UK speaking English and met the safety requirements to undergo magnetic resonance imaging (MRI).

Participants in the typically developing (TD) group presented with no history of speech and language problems and scored one deviation below the normative mean on no more than one standardized test score of language abilities. Participants in the DLD group presented with a history of speech and language problems and scored more than one standard deviation below the normative mean in two or more of the standardized language tests. Children who presented with a history of speech and language problems, including a previous DLD diagnosis, but failed to meet our DLD criteria at the time of testing, formed a separate group hereafter referred to as HSL.

### 2.2 Behavioral Measures

All participants who passed our first screening, completed a comprehensive neuropsychological test battery, providing a thorough and in-depth assessment of language and related abilities as well as motor coordination [see Supplementary Material Table 1 for a list of domains and tests used as part of the battery and see Krishnan et al., (2021) for a detailed description].

To reduce dimensionality and minimize multiple comparison problems, performance in the language and memory tests was summarized using factor scores. The approach used to identify the best weighted combination of these measures to give a language and a memory factor is described in Krishnan et al., (2021). In brief, a two-factor hybrid exploratory-confirmatory approach (E-CFA; Brown, 2015) was applied and compared against a single factor model. Model fit was significantly better (as indicated using Akaike’s information and Bayesian information criteria) for the two-factor compared to the one-factor model (see Krishnan et al., (2021) for a detailed description). Group mean and standard deviation for each factor (Language and Memory) and non-verbal IQ can be seen in Table (1).

### 2.3 MRI Acquisition

Imaging data were acquired using a 3T Siemens Prisma scanner with a 32-channel head coil. Participants wore noise-cancelling headphones (Optoacoustics OptoActive II Active Noise Cancelling Headphones), which were held in place using inflatable pads. Foam padding was inserted around the head for comfort and to restrict movement. High-resolution structural images were acquired using a MP-RAGE (Magnetization Prepared - Rapid Gradient Echo) T1-weighted sequence (TR = 1900 ms, TE = 3.97 ms, flip angle = 8°, FoV = 192 mm, 1 mm isotropic voxel size). Diffusion-weighted MRI followed the same sequence acquisition protocol used in the UK Biobank Project (Miller et al., 2016). In brief, sampling in *q*-space was performed in two shells at *b* = 1000 s/mm^2^ and 2000 s/mm^2^ (voxel size = 2 mm, MB factor = 3). For both diffusion-weighted shells, 50 distinct diffusion-encoding directions were acquired, covering 100 distinct directions across both shells. Five *b* = 0 s/mm^2^ images were obtained as well as three *b* = 0 s/mm^2^ images with reversed phase-encoding direction.

### 2.4 Imaging Data Analysis

#### 2.4.1 Pre-processing

Diffusion-weighted imaging data were processed using the FMRIB Software Library v6.0 (FSL; Smith et al., 2004). Prior to any pre-processing steps, all non-diffusion (b0) images for the anterior-posterior as well as the posterior-anterior phase-encoding direction were manually checked for artefacts. The first anterior-posterior and posterior-anterior b0 image was used as a default, however, in the presence of artefacts the best alternative b0 image was chosen. We started by estimating the susceptibility-induced off-resonance field from the pairs of images using the TOPUP correction tool (Andersson et al., 2003). We then performed skull stripping using FSL’s brain extraction tool (BET; Smith, 2002) creating a non-diffusion brain mask. Subsequently, we used FSL’s EDDY tool to correct diffusion-weighted imaging data for eddy current-induced distortions and participant head motion (Andersson & Sotiropoulos, 2016). We used the *mporder* option (Andersson et al., 2017) to correct for slice-to-volume (i.e. within-volume) movement as well as the *estimate_move_by_susceptibility* option (Andersson et al., 2018) to correct for susceptibility-by-movement interactions with eddy currents. Outlier detection was performed using the *repol* option (Andersson et al., 2016) to identify slices with signal loss due to motion during the diffusion encoding. Identified slices were replaced by non-parametric predictions using the Gaussian Process (Andersson et al., 2016). We further run automated quality control (EDDY QC) to detect data acquisition and pre-processing issues at subject and group level (Bastiani et al., 2019) [see Table 1 for Signal-to-Noise ratio (SNR) and relative motion displacement descriptives per group].

#### 2.4.2 Fitting the DWI data using tensor and NODDI models

After pre-processing, we fitted a tensor model at each voxel using b = 1000 s/mm^2^ sampled data (DTIFIT; Behrens et al., 2003) to compute fractional anisotropy (FA), eigenvalues (including axial diffusivity; AD) and eigenvector maps. The computed eigenvalues were then used to derive maps for radial diffusivity (RD).

Additional microstructural parameters were derived from neurite orientation dispersion and density imaging (NODDI; Zhang et al., 2012), using both shells of the diffusion data (1000 s/mm^2^ and 2000 s/mm^2^). NODDI protocols model diffusion data in three compartments: intra-cellular, extra-cellular, and cerebrospinal fluid (CSF). NODDI parameters were estimated with the CUDA diffusion modelling toolbox (cuDIMOT; Hernandez-Fernandez et al., 2019) using the Watson model with Markov Chain Monte Carlo optimization. Three parameters were estimated: fraction of the data that is isotropic (f_iso;_ e.g., how much of a voxel is associated with CSF); fraction of intra-cellular compartment compared to the total intra- and extra-cellular compartment (f_intra_; that is, disregarding f_iso_); and orientation dispersion (OD). OD provides a measure of how disperse the fibers are, bounded from 0 to 1. A low OD indicates that the fibers are aligned; high OD means fiber directions are not aligned. Changes in OD and f_intra_ both impact FA, as would partial volume effects of CSF captured by f_iso_. NODDI parameters may help disentangle the microstructural differences underlying differences in FA and MD (Zhang et al., 2012).

#### 2.4.3 Tractography

We estimated the probability distribution of diffusion directions using BEDPOSTX (Jbabdi et al., 2012). The transformation from diffusion to standard space (and vice versa) was calculated using non-linear registration with the subject-specific T1-weighted anatomical image as an intermediate image in the transformation (FNIRT; Andersson et al., 2010). Probabilistic tractography was performed using XTRACT (Warrington et al., 2020), which transforms standard space masks for seeds and targets into each participant’s native diffusion space. The inferior (ICP) and superior cerebellar peduncles (SCP) in each hemisphere as well as the middle cerebellar peduncle (MCP) were identified using a published protocol and available masks by Bruckert et al., (2019). In brief, masks were used as seeds and targets in XTRACT to robustly segment portions of the peduncles. For the left and right ICP, the seed mask was a region-of-interest (ROI) placed in an axial slice at the level of the pontomesencephalic junction and the target was a second ROI placed ipsilaterally in an axial slice at the level of the medulla oblongata (target mask). The MCP was segmented by placing ROIs in the central portion of the left (seed mask) and right (target mask) MCP in an axial plane at the medial pons level. The left and right SCP were segmented by placing the seed mask in the SCP in an axial plane at the level of the pontomesencephalic junction and the target mask in the dentate nucleus in an axial plane at medial pons level. We added a midline exclusion mask to prevent the streamlines crossing and the seed and target masks were used as stop/termination masks to prevent streamlines continuing past these ROIs. Note that the midline exclusion mask for the MCP allowed pontine streamlines to cross to the contralateral hemisphere. Figure 1 shows the masks for the ICP (A), MCP (B), and the SCP (C) from the coronal, axial, and sagittal perspective. We used XTRACT default settings; the resulting tract segments of the peduncles were thresholded to remove the bottom 10%, binarized, and used to mask the diffusion images. Figure 2 displays the tractography output in a representative TD and DLD child (see Supplementary Material Figure S2 for a representative HSL child). Means for each tract for FA, AD, RD, OD, f_intra_, f_iso_ were obtained along with whole-brain FA for each participant.

**Figure 1:**
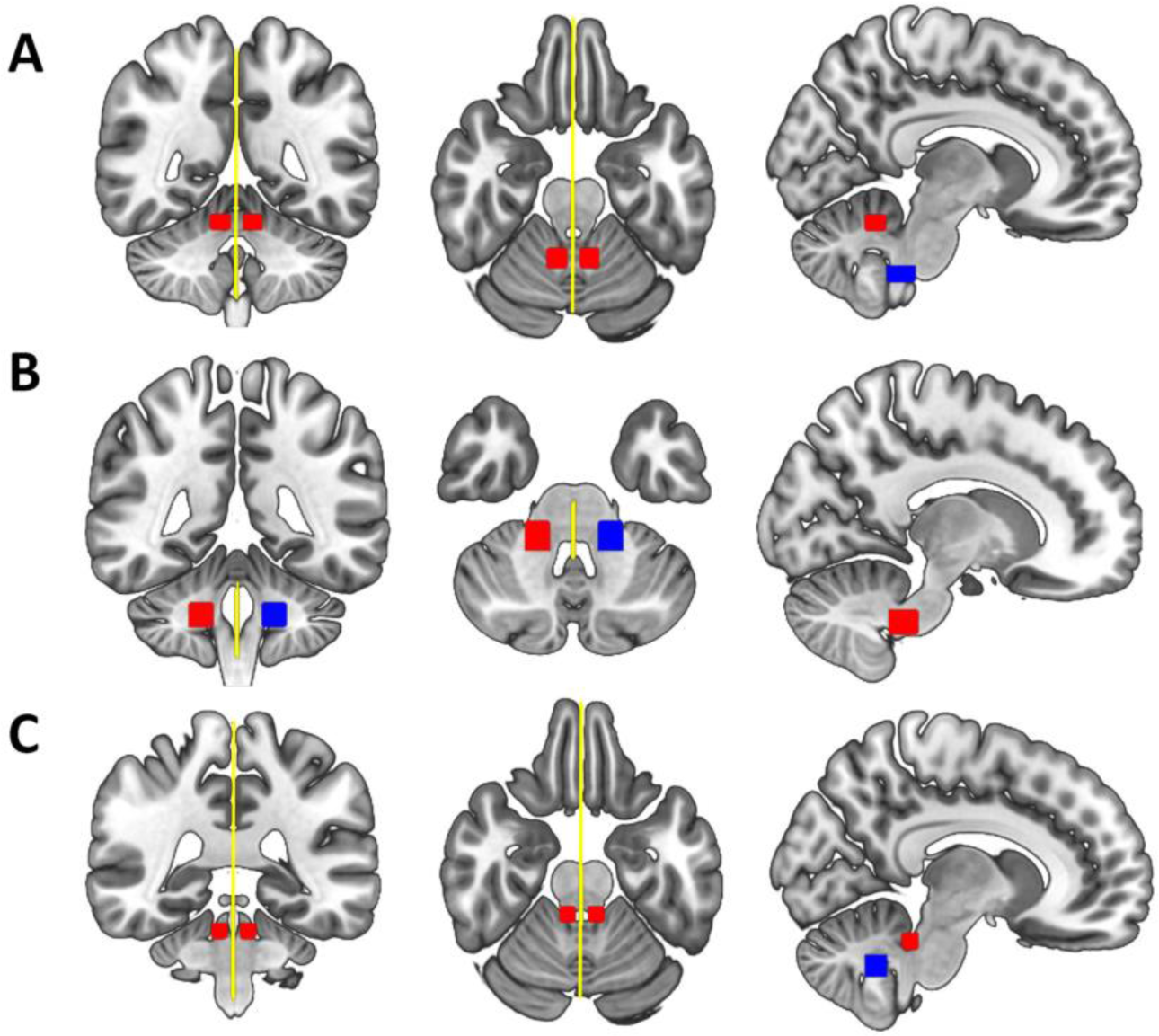
Delineation of the Bruckert et al. (2019) masks used to extract (A) the left and right inferior cerebellar peduncles, (B) the middle cerebellar peduncle, and (C) the left and right superior cerebellar peduncles from the coronal (left), axial (middle) and sagittal (right) perspective displayed in standard space. Seed masks are colored in red, target masks in blue, and exclusion masks in yellow. An additional stop mask was placed at seed positions (not displayed in the figure).

**Figure 2:**
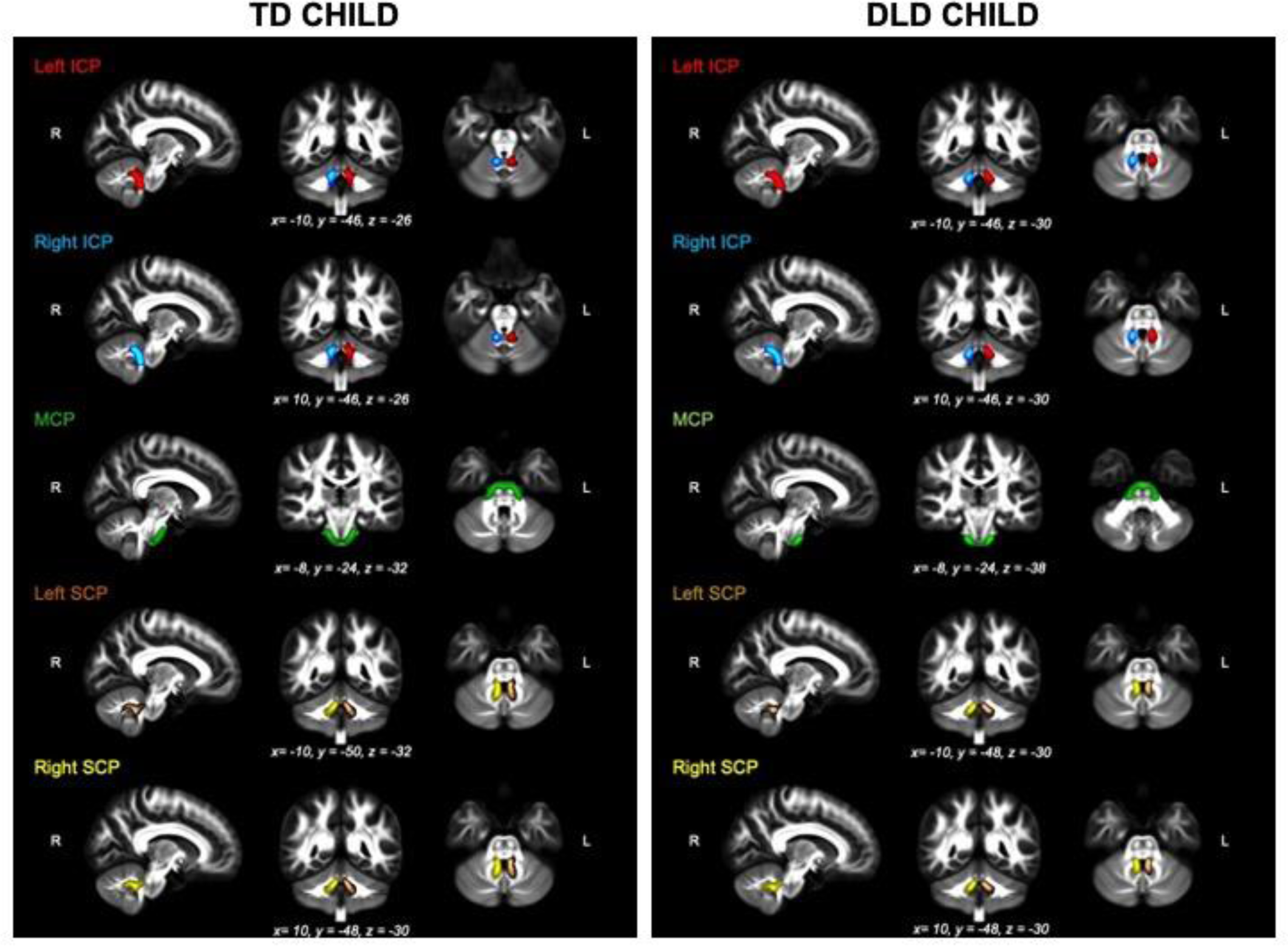
The thresholded binarized inferior (ICP; left in red, right in blue), middle (MCP; green) and Superior (SCP; left in brown, right in yellow) cerebellar peduncles overlaid on the FSL_HCP065_FA image in a typical TD (left panel) and DLD child (right panel).

#### 2.4.4 Cerebellar Volume Analysis

We used Freesurfer (version 7.2.0; http://surfer.nmr.mgh.harvard.edu/) to perform volumetric segmentation of participants’ T1-weighted images (Dale et al., 1999). Preprocessing included skull-stripping, automated Talairach transformation, subcortical segmentation of gray and white matter volumetric structures (including the cerebellum), and automatic labeling of brain volume (Fischl et al., 2002, 2004). Data from three TD participants were excluded after quality control. Participants’ left and right cerebellar grey and white matter volumes (measured in mm^3^) were obtained from the segmentation output for statistical group comparisons. Variation in whole brain volume was corrected for by statistically regressing out the Freesurfer estimated intracranial volume (ICV) for each participant. A linear mixed effects model (lme4 package, Bates et al., 2014) was run with volume (in mm^3^) as the dependent variable, and group (TD, DLD), hemisphere (left, right), tissue (grey, white), group-by-hemisphere-by-tissue interaction, and total intracranial volume as predictors (formula = lmer(volume ∼ group*hemisphere*tissue + ICV + (1|sub), Data).

### 2.5 Statistical Analysis

All statistical analyses were conducted in R version 4.2.1 (R Core Team, 2022). We analyzed group differences in FA using generalized linear mixed effects analysis modelled using beta distribution as implemented in the *glmmTMB* package (Brooks, Mollie et al., 2017). Beta regression models, as introduced by Ferrari & Cribari-Neto, (2010), are used when the dependent variable is beta distributed, i.e., continuous probability distributions defined on the interval [0,1]. This is suited for FA data that range from 0 (isotropic) to 1 (completely anisotropic).

Participants’ extracted mean FA values for each of the peduncles (ICP, MCP, SCP) were the dependent variable in the generalized linear mixed models. As fixed effects we entered group (TD, DLD, HSL), hemisphere (left and right in the ICP and SCP) and their interaction, as well as whole-brain FA as a covariate. As random effects we entered intercepts for participants (1|subj). We then adjusted each of the models for age, sex, and relative motion displacement during scan acquisition, to assess whether any significant effects would remain significant in the adjusted models.

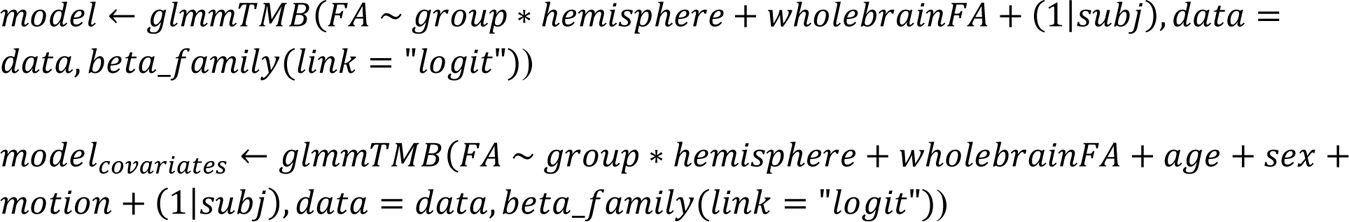

We used Bonferroni-corrected alpha levels based on the number of models tested to adjust for multiple comparisons where appropriate.

## 3. Results

Probabilistic tractography was successful in all 159 participants. We reviewed imaging quality and tractography results for each participant by visually inspecting registration, DTIFIT, NODDI, and tractography output. Data quality was deemed sufficient for all participants and no observations were excluded from further analyses. A table summarizing mean FA in each peduncle for the three groups can be found in Supplementary Material Table (S3).

### 3.1 Group effects on FA in Cerebellar Peduncles

We tested whether FA in the cerebellar peduncles can be predicted by group (TD, DLD, HSL) by fitting three beta-distributed generalized linear mixed effects models – one per peduncle. To adjust for multiple comparisons, we applied a Bonferroni-corrected alpha level of 0.017.

#### 3.1.1 Inferior Cerebellar Peduncle

The model predicting mean FA in the ICP (formula = glmmTMB(FA ∼ group*hemisphere + whole-brain FA + (1|subj), Data, beta_family(link = “logit”)) showed a significant main effect of group (*b* = −0.09, *SE* = 0.04, 95% CI = [-0.17,-0.02], *z* = −2.48, *p_uncorrected_* = 0.013). Mean FA values in the ICP were lower in the DLD group (Left ICP *M* = 0.42, *SD* = 0.06; Right ICP *M* = 0.45, *SD* = 0.05) compared with the TD group (Left ICP *M* = 0.45, *SD* = 0.06; Right ICP *M* = 0.47, *SD* = 0.04). This effect remained significant after adjusting for age, sex, and head motion. FA data are presented in Figure 3 and the full model summaries are available in Table 2. These results remain the same if we exclude the HSL group from the analysis (see Supplementary Material Table S4).

**Figure 3:**
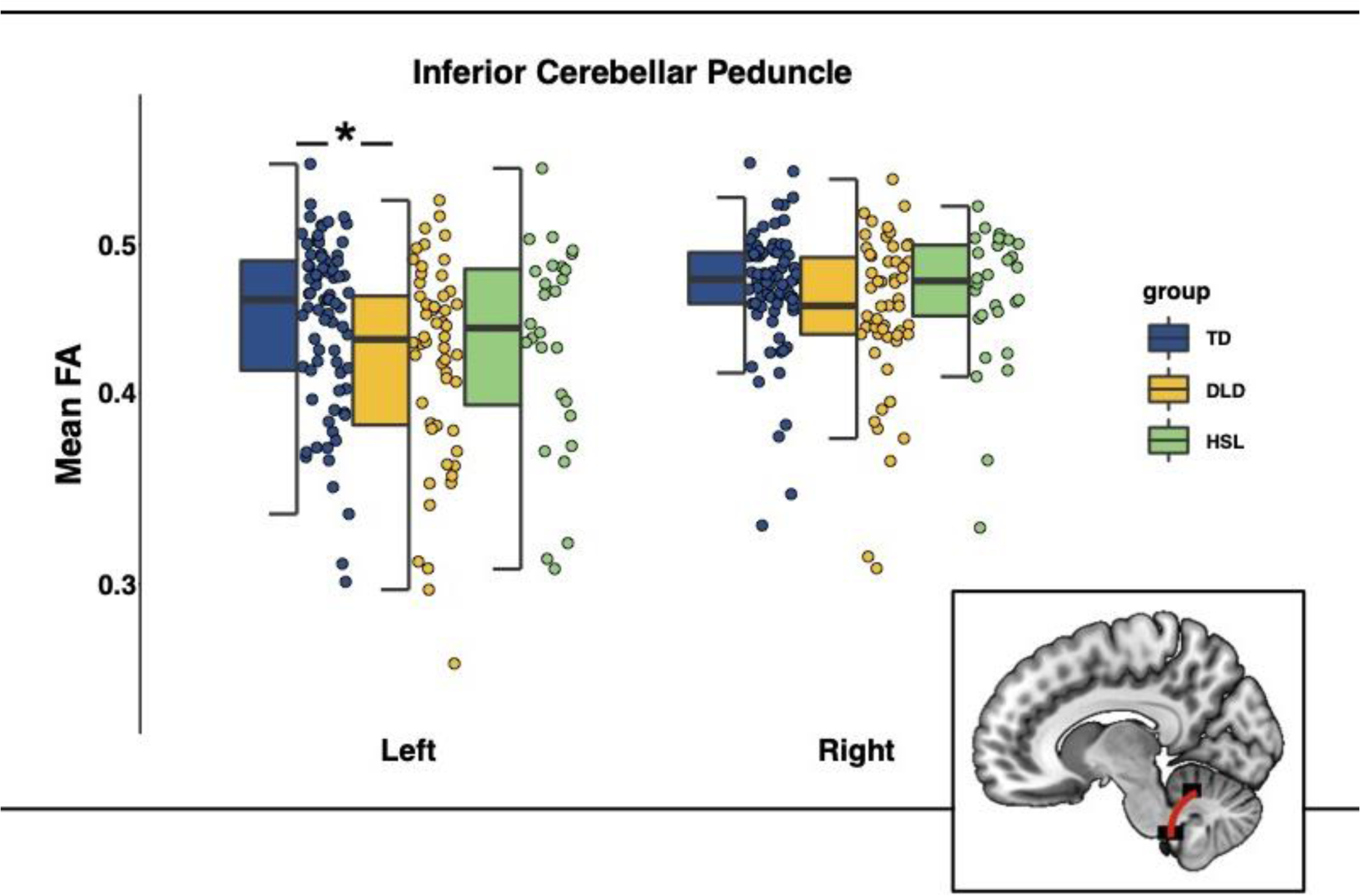
Mean Fractional anisotropy (FA) in the inferior cerebellar peduncles (ICP) by group (TD = typically developing in blue, DLD = developmental language disorder in yellow, HSL = history of speech and language impairments in green) and across hemispheres. Graphical representation of ICP in red overlaid on a standardized template in sagittal view. *Note:* The box is drawn from first to third quartile with the horizontal line drawn in the middle to denote median FA. Whiskers show 1.5*IQR with data beyond the end of the whiskers (group outliers) being plotted individually. The asterisk denotes the significant difference in FA between TD and DLD in the left ICP, as indicated by post-hoc pairwise comparisons (Bonferroni corrected).

**Table 2:** Model summaries for fractional anisotropy (FA) in the inferior cerebellar peduncles (ICP).

| Predictor | ICP Model |  |  | ICP Model including Age, Sex, Motion |  |  |
| --- | --- | --- | --- | --- | --- | --- |
|  | Coefficient | 95% CI | p-value | Coefficient | 95% CI | p-value |
| Group |  |  |  |  |  |  |
| TD | — | — |  | — | — |  |
| DLD | -0.09 | -0.17, -0.02 | 0.013* | -0.08 | -0.15, -0.01 | 0.031 |
| HSL | -0.06 | -0.15, 0.03 | 0.2 | -0.03 | -0.12, 0.06 | 0.5 |
| Hemisphere |  |  |  |  |  |  |
| Left | — | — |  | — | — |  |
| Right | 0.09 | 0.04, 0.14 | 0.001* | 0.09 | 0.04, 0.14 | 0.001 |
| Whole brain FA | 1.2 | -2.1, 4.6 | 0.5 | 1.5 | -1.8, 4.8 | 0.4 |
| Group * Hemisphere |  |  |  |  |  |  |
| DLD * r | 0.03 | -0.05, 0.12 | 0.4 | 0.03 | -0.05, 0.12 | 0.4 |
| HSL * r | 0.03 | -0.08, 0.13 | 0.6 | 0.03 | -0.08, 0.13 | 0.6 |
| Age |  |  |  | 0.00 | -0.02, 0.02 | 0.9 |
| Sex |  |  |  |  |  |  |
| Male |  |  |  | — | — |  |
| Female |  |  |  | 0.10 | 0.04, 0.16 | <0.001 |
| Head motion |  |  |  | 0.09 | -0.14, 0.32 | 0.5 |

Note: TD (typically developing) group acts as reference category for the other groups (DLD = developmental language disorder, HSL = history of speech and language impairments). CI = Confidence Interval; Coefficient = unstandardized (b). An asterisk (\*) indicates p-values that survived Bonferroni correction.

Post-hoc pairwise comparisons revealed a significant difference between TD and DLD in FA in the left ICP [*b* = 0.09, *SE* = 0.04, *t*(309) = 2.48, *p_Bonferroni_* = 0.041].

To gain a better understanding of the microstructure underlying the FA differences in the ICP, we examined mean radial diffusivity (RD), diffusivity perpendicular to the axonal tract which is often used as an index of myelination, and axial diffusivity (AD), diffusivity parallel to the tract which is often used as an index of axonal integrity. FA is a ratio of AD to RD, so changes in either RD or AD (or both) can affect FA estimates. The model predicting mean RD in the ICP (formula = glmmTMB(RD ∼ group*hemisphere + whole-brain FA + (1|subj), Data)) revealed no differences among groups. The model predicting mean AD in the ICP (formula = glmmTMB(AD ∼ group*hemisphere + whole-brain FA + (1|subj), Data)) revealed a significant main effect of group (DLD group *b* = −2.79×10^-5^, *SE* = 9.64×10^-6^, 95% CI [-4.66×10^-5^,-9.13×10^-6^], *t* = −2.89, *p_uncorrected_* = 0.004) such that participants in the DLD group had lower AD (Left ICP *M* = 1.16×10^-3^, *SD* = 5.52×10^-5^; Right ICP *M* = 1.18×10^-3^, *SD* = 5.31×10^-5^) compared with participants in the TD group (Left ICP *M* = 1.19×10^-3^, *SD* = 6.00×10^-5^; Right ICP *M* = 1.19×10^-3^, *SD* = 5.31×10^-5^). Post-hoc pairwise comparisons revealed a significant difference between TD and DLD in AD in the left ICP [TD – DLD: *b* = 2.79 × 10^-5^, *SE* = 9.64 × 10^-6^, *t*(269) = 2.89, *p_Bonferroni_* = 0.013].

We also fitted our diffusion data using the NODDI tissue model which allowed us to compare OD, f_intra_, and f_iso_ in the ICP among groups. These measures reflect microstructural complexity of dendrites and axons which contributes to diffusion tensor indices such as FA. No differences among groups were found for FINTRA (formula = glmmTMB(FINTRA ∼ group*hemisphere + whole-brain FA + (1|subj), Data, beta_family(link = “logit”))

or FISO (formula = glmmTMB(FISO ∼ group*hemisphere + whole-brain FA + (1|subj), Data, beta_family(link = “logit”)). We did, however, find a significant group effect in the model predicting mean OD in the ICP (formula = glmmTMB(OD ∼ group*hemisphere + whole-brain FA + (1|subj), Data, beta_family(link = “logit”)) DLD group *b* = 0.11, *SE* = 0.04, 95% CI [0.047, 0.182], z = 3.32, *p_uncorrected_* < 0.001. Participants in the DLD group had higher OD (Left ICP *M* = 0.27, *SD* = 0.04; Right ICP *M* = 0.24, *SD* = 0.04) compared with participants in the TD group (Left ICP *M* = 0.24, *SD* = 0.04; Right ICP *M* = 0.23, *SD* = 0.03). Post-hoc pairwise comparisons revealed a significant difference between TD and DLD in OD in the left ICP [TD – DLD: *b* = - 0.12, *SE* = 0.04, *t*(309) = −3.32, *p_Bonferroni_* = 0.003].

#### 3.1.2 Middle & Superior Cerebellar Peduncles

The model predicting mean FA in the MCP (formula = glmmTMB(FA ∼ group + whole-brain FA + (1|subj), Data, beta_family(link = “logit”))) showed no effect of group (DLD group: *b* = - 0.01, *SE* = 0.03, 95% CI [-0.06,0.04], *z* = −0.42, *p* = 0.68; HSL group: : *b* = −0.03, *SE* = 0.03, 95% CI [-0.09,0.04], *z* = −0.89, *p* = 0.38; TD group: reference group). Similarly, no effect of group was found in the model predicting mean FA in the SCP (formula = glmmTMB(FA ∼ group*hemisphere + whole-brain FA + (1|subj), Data, beta_family(link = “logit”))) (DLD group: *b* = −0.04, *SE* = 0.02, 95% CI [-0.08, 0.01], *z* = −1.50, *p* = 0.13; HSL group: : *b* = 0.05, *SE* = 0.03, 95% CI [-0.10, 0.10], *z* = 1.60, *p* = 0.11; TD group: reference group) (see Figure 4).

**Figure 4:**
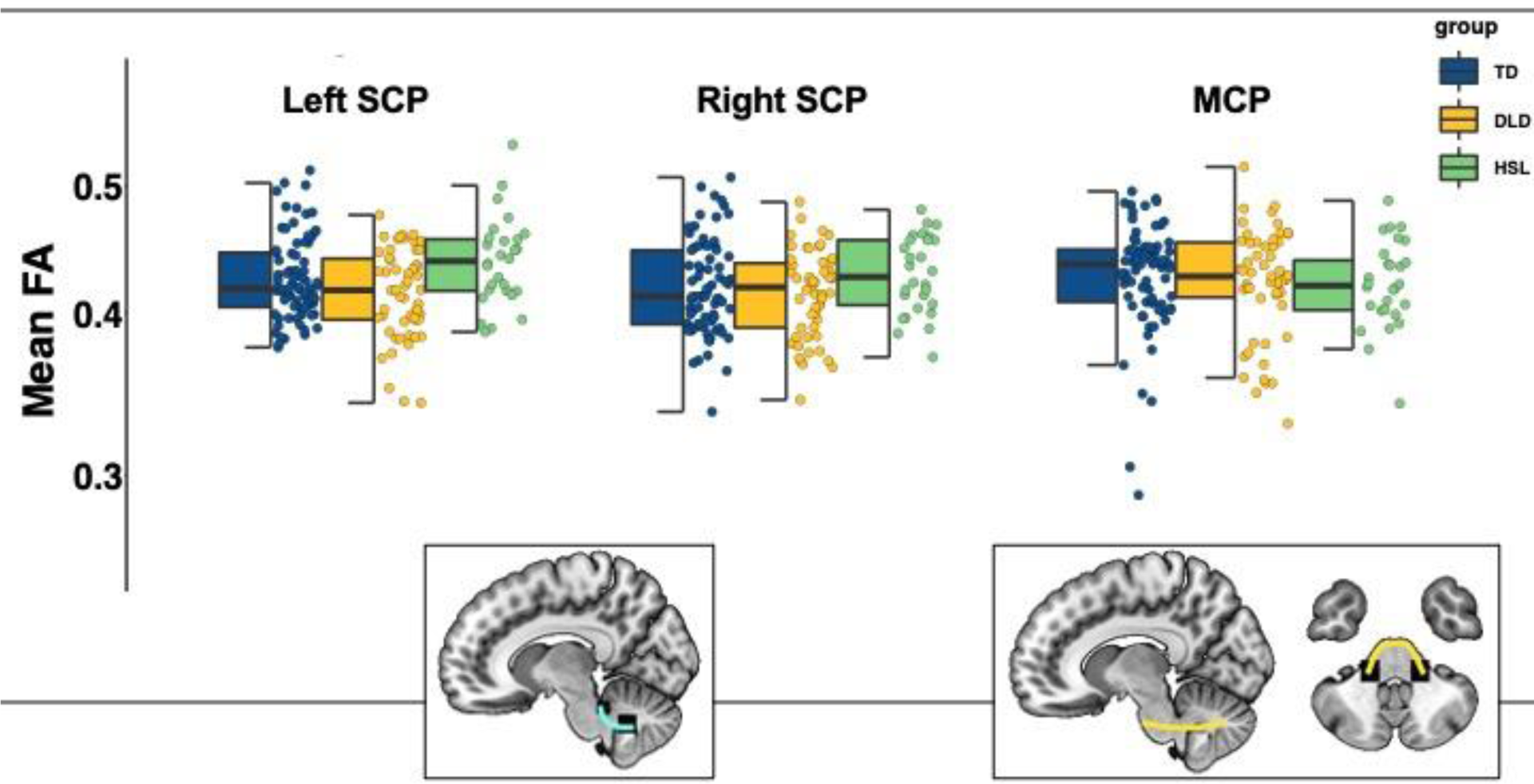
Mean Fractional anisotropy (FA) in the superior and middle cerebellar peduncles by group (TD = typically developing in blue, DLD = developmental language disorder in yellow, HSL = history of speech and language impairments in green) and across hemispheres. Graphical representations of SCP (cyan) and MCP (yellow) overlaid on a standardized template in sagittal and axial view. *Note:* The box is drawn from first to third quartile with the horizontal line drawn in the middle to denote median FA. Whiskers show 1.5*IQR with data beyond the end of the whiskers (group outliers) being plotted individually. There were no significant differences among groups in these tracts.

### 3.2 Associations between behavioral outcomes and FA in the ICP

Having identified group differences in the inferior cerebellar peduncles using a categorical approach, we tested for associations between FA in these tracts and behavioral outcome measures including language, memory, and motor coordination. In this continuous analysis approach, we fitted linear and linear multivariate regression models predicting behavioral performance from FA in the left and right ICP across groups. We included whole-brain FA as a control covariate in these models (formula = lm(cbind(memory, language) ∼ FA + whole-brain FA, Data; formula = lm(pegboard ∼ FA + whole-brain FA, Data). To adjust for multiple comparisons, we applied a Bonferroni-corrected alpha level of 0.0125 to all 4 regression models. We found no statistically significant relationship between any of the behavioral measures and FA in the left and right ICP.

### 3.3 Testing for Group by Age Interaction effects

A previous study by Lee and colleagues (2020) tested FA in the MCP and SCP in adolescent and young adults with DLD (14 - 28 yrs) and found significant group and group-by-age interaction effects. The interaction revealed distinct developmental trajectories in DLD vs. TD participants: while FA increased with age in TD participants it decreased in DLD participants (Lee et al., 2020). We decided to run the same analysis on our sample, to see whether the findings would replicate in a younger but overlapping age-range. We ran beta-distributed generalized linear mixed effects models for each tract separately with group (TD, DLD), age, gender, handedness, and non-verbal IQ as fixed effects, and intercepts for participants as random factor (formula = glmmTMB(FA ∼ group*age + gender + handedness + non-verbal IQ + (1|subj), Data, beta_family(link=”logit”)). We further expanded this model to the left and right ICP which were not tested in Lee et al. (2020). To adjust for multiple tests (five regressions), we applied a Bonferroni-corrected alpha level of 0.01.

No significant group or group-by-age interaction effects were found in FA in the MCP, SCP (left and right). There was, however, a significant main effect of group (*b* = 0.62, *SE* = 0.22, 95% CI [0.19,1.04], *z* = 2.82, *p_uncorrected_* = 0.005) and a significant age*group interaction effect (*b* = - 0.06, *SE* = 0.02, 95% CI [-0.001,-0.02], *z* = −3.16, *p_uncorrected_* = 0.002) in the right ICP (see Figure 5). Similar effects were found in the left ICP but they did not survive correction. Full ICP model summaries can be found in Table 3.

**Figure 5:**
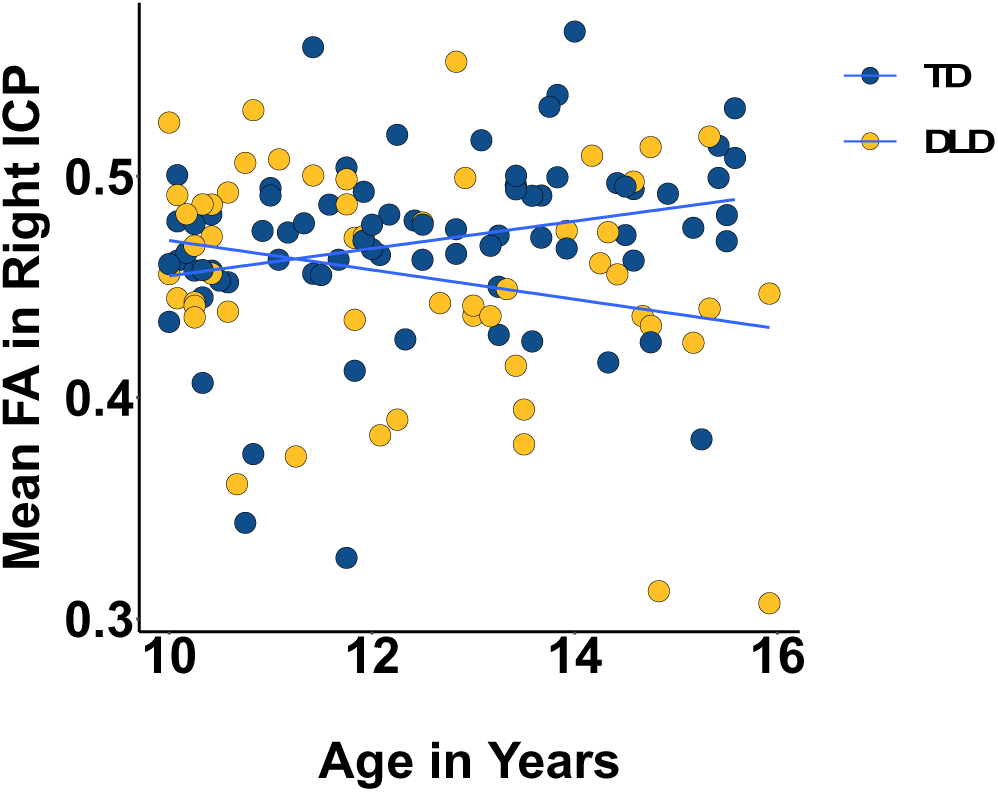
Mean Fractional Anisotropy (FA) in the right (bottom) inferior cerebellar peduncle by age and group (TD = typically developing in blue, DLD = developmental language disorder in yellow). The slopes were significantly different from zero and also significantly different from each other.

**Table 3:** Model summary for fractional anisotropy (FA) of the fiber tracts in the inferior (ICP), middle (MCP), and superior cerebellar peduncles (SCP) as tested by Lee et al. (2020) using group (TD vs. DLD), age, sex, handedness, and non-verbal IQ as fixed effects.

| Predictor | ICP Left |  |  | ICP Right |  |  |
| --- | --- | --- | --- | --- | --- | --- |
|  | Beta | 95% CI<sup>1</sup> | p-value | Beta | 95% CI<sup>1</sup> | p-value |
| Group |  |  |  |  |  |  |
| TD | — | — |  | — | — |  |
| DLD | 0.52 | -0.03, 1.1 | 0.066 | 0.61 | 0.19, 1.0 | 0.005* |
| Age | 0.02 | -0.01, 0.05 | 0.11 | 0.03 | 0.00, 0.05 | 0.030 |
| Sex |  |  |  |  |  |  |
| Male | — | — |  | — | — |  |
| Female | 0.09 | 0.01, 0.18 | 0.029 | 0.06 | 0.00, 0.13 | 0.061 |
| Handedness |  |  |  |  |  |  |
| Right | — | — |  | — | — |  |
| Left | -0.06 | -0.17, 0.05 | 0.3 | 0.02 | -0.06, 0.10 | 0.7 |
| Non-verbal IQ | 0.01 | -0.01, 0.02 | 0.5 | 0.00 | -0.02, 0.01 | 0.5 |
| Group * Age |  |  |  |  |  |  |
| DLD * Age | -0.05 | -0.09, 0.00 | 0.040 | -0.05 | -0.09, -0.02 | 0.002* |

*Note: TD (typically developing) group acts as reference category for the other group (DLD = developmental language disorder). CI = Confidence Interval; Coefficient = unstandardized (b). An asterisk (\*) indicates p-values that survived Bonferroni correction.*

The significant interaction was probed using the *emmeans* package (Lenth, 2023) to assess the statistical significance of simple slopes against zero and simple slope differences for age across the two groups. The slope of age on FA in the TD group was 0.025, 95% CI [0.002, 0.048] while in DLD it was –0.03, 95% CI [-0.056, −0.004]. The 95% confidence interval did not contain zero for either group, so the simple slope was significant for both. The difference in slopes was also significant [TD – DLD: *b* = 0.06, *SE* = 0.02, *t*(121) = 3.16, *p* = .002] with TD having a higher slope than DLD.

### 3.4 Volumetric differences in the cerebellum

We also examined differences in cerebellar grey and white matter in DLD, using Freesurfer anatomical segmentation output. There was a significant effect of group (*b* = −1564, *SE* = 733.3, *t*(353.2) = −2.13, *p* = .034) such that the DLD group (Left Cerebellum: Cortex M = 56317.72, SD = 6126.39, White Matter : M = 13683.67, SD = 1836.93; Right Cerebellum Cortex M = 56864.42, SD = 6503.32, White Matter : M = 113051.03, SD = 1663.48) had smaller cerebellar volume than the TD group (Left Cerebellum: Cortex M = 58072.65, SD = 5503.35, White Matter : M = 14838.51, SD = 1643.63; Right Cerebellum Cortex M = 58767.31, SD = 5710, White Matter : M = 13997.30, SD = 1528.11). The three-way interaction was not significant. Post-hoc pairwise comparisons indicated differences in the left [*b* = 1564, *t*(353) = 2.13, *p_uncorrected_* = 0.034] and right cerebellar cortex [*b* = 1721, *t*(353) = 2.35, *p_uncorrected_* = 0.02], however these did not survive Bonferroni correction for multiple comparisons.

## 4. Discussion

Cortico-cerebellar circuits, alongside cortico-striatal systems, contribute to complex sensorimotor skill learning, including the acquisition of spoken language (Stoodley, 2016; Ziegler & Ackermann, 2017). In this study, we hypothesized that abnormalities in cortico-cerebellar circuits are associated with language learning problems in development. To this end, we investigated differences in cerebellar white matter connectivity in children with DLD, a common developmental language disorder, using diffusion weighted imaging. We found that FA in the left ICP was lower in children with DLD, and that this was primarily driven by decreased AD and increased OD in DLD compared to TD children. We also found atypical age-related changes in the right ICP such that while FA increased with age in TD children it decreased in DLD, put another way, the group difference in FA in the right ICP is only evident at later ages and FA values in the two groups are overlapping at the younger end of the age range. We did not, however, find any associations between FA in the ICP and language, memory, or motor performance. These findings indicate typical cortico-cerebellar (MCP and SCP) but atypical olivocerebellar connectivity in DLD and suggest that the olivocerebellar system might play in important role in speech development.

Considering previous research findings in DLD, which have implicated atypical perisylvian and striatal structure and function, as well as the broad range of difficulties that children with DLD present with (expressive and receptive language, vocabulary, grammar, narrative, etc.), we anticipated differences primarily in the cortico-cerebellar pathways (i.e., the middle and superior cerebellar peduncles). Instead, we identified differences in the ICP, a pathway that feeds the cerebellum with sensory inputs (including proprioceptive and visual inputs) from neurons located in the periphery, including somatosensory input from speech organs (oral and facial muscles) (Moberget & Ivry, 2016). It has been proposed that this input regarding the current sensory state is combined with feed-forward input from the cortex to allow computation of predictions (or internal models) and prediction-errors (Moberget & Ivry, 2016). These cerebellar computations ultimately contribute to performance refinement and learning via error correction (Caligiore et al., 2016; Koziol et al., 2014).

Animal studies have highlighted the role of the olivocerebellar pathway in motor learning and motor control (Lang et al., 2016) and are corroborated by patient studies that show that lesion in this pathway impaired performance in motor adaptation tasks (Martin et al., 1996). More recently, a study reported that motor adaptation performance in neurotypical controls was negatively correlated with FA in the left ICP (Jossinger et al., 2020) offering more evidence that adaptation relies on afferent cerebellar input. With respect to speech and language disorders, the ICP has been repeatedly implicated in developmental stuttering, a common developmental disorder characterized by speech dysfluency. Fractional anisotropy was reduced in the ICP as well as the other cerebellar peduncles in adults who stutter compared to controls (Connally et al., 2014). Reduced FA in the right ICP, but not in other peduncles, was found in children who stutter, and FA in this tract showed a negative relationship with stuttering frequency (Johnson et al., 2022). Lastly, a negative association was found between speech rate and FA in the left ICP in adults who stutter, potentially facilitating a hyperactive error-correction process from afferent inputs and resulting in lower speech rates in this group (Jossinger et al., 2021).

Only a handful of studies have considered the role of the cerebellum in DLD – perhaps unsurprisingly given the scarcity of neuroimaging studies on DLD. One recent study reported larger grey matter volume in the right cerebellum in children with DLD compared with TD – an unanticipated finding as the authors hypothesized (and did not find) differences in the caudate nucleus, the IFG and STG instead (Pigdon et al., 2019). This is in agreement with our current findings and the broader literature as atypical cerebellar volume has also been reported in a number of developmental disorders such as ASD, ADHD, and dyslexia (Stoodley, 2016). With respect to cerebellar connectivity in DLD, one recent study reported lower FA in the SCP and MCP bilaterally (the ICP was not tested) in DLD – this was again an unforeseen finding as the authors anticipated differences in cortico-striatal tracts (Lee et al., 2020). The discrepancy between Lee et al.’s (2020) findings and ours could be due to differences in age (their sample consisted of adolescents and young adults with DLD), sample size, as well as imaging and analysis protocols.

Previous findings on the SCP and MCP in DLD emphasized the importance of age as a moderator when considering group differences (Lee et al., 2020). Indeed, we found that while FA increased with age in TD children, it did not in children with DLD in the right ICP (with the left ICP following a similar trend). Longitudinal studies have shown that the middle and superior cerebellar peduncles reach their white matter maturation peak between 12-15 years, while the ICP between 12-17 years (Simmonds et al., 2014). The age range of our sample (10-16 years) was therefore well-suited for investigating the developmental trajectories of these tracts, indicating an atypical time course of maturation in the ICP in children with language learning problems. Alternatively, it could be the case that older children with DLD were simply more “severe” in their language profile, a possibility we cannot exclude in our cross-sectional approach.

Fractional anisotropy is sensitive to white matter microstructural properties, including myelin content, however differences in FA do not necessarily translate to differences in myelination (Jones et al., 2013). The group differences we observed in FA in this study are driven by reduced axial diffusivity and increased orientation dispersion in DLD. This suggests that reduced orientation coherence of neurites is driving the effect rather than radial diffusivity and neurite density, through myelination and axon density (Jespersen et al., 2010). Unlike FA, OD is relatively stable across development with only a weak (negative) correlation with age (Lynch et al., 2020). A recent study has also shown that OD in the peduncles is highly heritable (Luo et al., 2022). Our results may therefore reflect causal factors at play in DLD, rather than a consequence of learning deficits.

The main limitation of the present study is the cross-sectional design which does not allow us to disentangle whether the differences we’ve observed are the cause or consequence of DLD. Future longitudinal studies are required to test the developmental trajectories of white matter pathways in DLD and to make any strong developmental claims. Another important limitation is that we used DWI tractography to identify and quantify the cerebellum’s white matter connectivity *in vivo*. While its utility is undeniable, tractography is inherently limited and can only offer an indirect and crude approximation of the anatomical fiber pathways (Jeurissen et al., 2019; Schilling et al., 2019; Thomas et al., 2014).

To conclude, we found lower FA in the ICP, the white matter pathway that carries sensory input from the periphery to the cerebellum, in children with DLD. Our results point to a potentially suboptimal transfer of sensory input in individuals with profound language-learning difficulties. Unlike cortical and striatal mechanisms, cerebellar contributions to learning might be more important early on during skill acquisition (Stoodley, 2016; Ziegler & Ackermann, 2017). According to this view, the information carried, and computations facilitated by the ICP might be selectively consequential for motor learning early on in speech acquisition. Alternatively, the ICP might support motor speech functions across the lifespan, a view partially supported by findings from non-invasive stimulation studies on the cerebellum which introduce disruptions to different motor but also speech and language functions in neurotypical adults (Arasanz et al., 2012; Lesage et al., 2012; Runnqvist et al., 2016). The two views are not mutually exclusive but in order to understand whether the cerebellum selectively affects typical and atypical language development more longitudinal data are needed. Finally, future studies should adopt a network approach to investigate the interactions between the cerebellar and the cortico-striatal systems known to be atypical in DLD. Such an approach may be crucial in understanding the neurobiological underpinnings of this complex disorder.

## Acknowledgements

We would like to acknowledge the efforts of the children and families who participated and gave up their personal time to help us learn about brain and language development. We gratefully acknowledge everyone who assisted us with participant recruitment (https://boldstudy.wordpress.com/acknowledgements/). We also wish to thank the radiographers at the Oxford Centre for Human Brain Activity: Nicky Aikin, Nicola Filippini, Emily Hinson, Sebastian Rieger, Juliet Semple, and Eniko Zsoldos. We would like to thank Dr Paul Thompson for providing statistical advice; Dr Amy Howard for advice on NODDI fitting; Nilgoun Bahar for performing the FreeSurfer quality control; and Prof Dorothy Bishop for providing guidance and feedback throughout this project.

## Data and Code Availability Statements

The code and anonymized data will be made openly available upon acceptance of the manuscript.

