## Supplementary Material for "Microstructural Properties of the Cerebellar Peduncles in Children with Developmental Language Disorder"

*Supplementary Table (S1)*: List of tests used as part of the neuropsychological battery. The measures under the *Language* and *Memory* domains were summarized into two factors using factor analysis.

| Domain | Skill | Test |
|---|---|---|
| Language | Receptive Grammar | Test for Reception of Grammar TROG-2; Bishop, 2003 |
| | Expressive Grammar | Clinical Evaluation of Language Fundamentals CELF-4 Sentence recall; Semel et al., 2004 |
| | Receptive Vocabulary | Receptive One-Word Picture Vocabulary Test ROWPVT-4; Martin and Brownell, 2011 |
| | Expressive Vocabulary | Expressive One-Word Picture Vocabulary Test EOWPVT-4; Martin and Brownell, 2011 |
| | Narrative Production & Comprehension | Expression, Reception and Recall of Narrative Instrument ERNNI; Bishop, 2004 |
| | Phonological Processing | Nonword repetition; Snowling et al., 2015 |
| Reading | Decoding & Word Reading | Test Of Word Reading Efficiency TOWRE; Torgesen et al., 1999 |
| Memory | Short-term & Working Memory | Forward and Backward Digit Span Children's Memory Scale CMS; Cohen, 1997 |
| | Episodic Auditory-Verbal Learning | Word lists CMS; Cohen, 1997 |
| Motor | Oromotor Coordination | Oromotor sequences subtest of the NEPSY (A Developmental NEuroPSYchological assessment); Korkman et al. 1998 |
| | Gross and fine Motor Dexterity | Purdue Pegboard; Tiffin, 1968 |
| Non-verbal IQ | Visuospatial Ability | Block Design Wechsler Intelligence Scale for Children WISC IV; Wechsler, 2004 |
| | Visual/abstract/perceptual Reasoning | Matrix Reasoning WISC IV; Wechsler, 2004 |
| | Processing speed | Coding WISC IV; Wechsler, 2004 |

*Supplementary Material Figure (S2)*: The thresholded inferior (ICP; left in red, right in blue), middle (MCP; green) and Superior (SCP; left in brown, right in yellow) cerebellar peduncles overlaid on the FSL_HCP065_FA image in a typical HSL child.

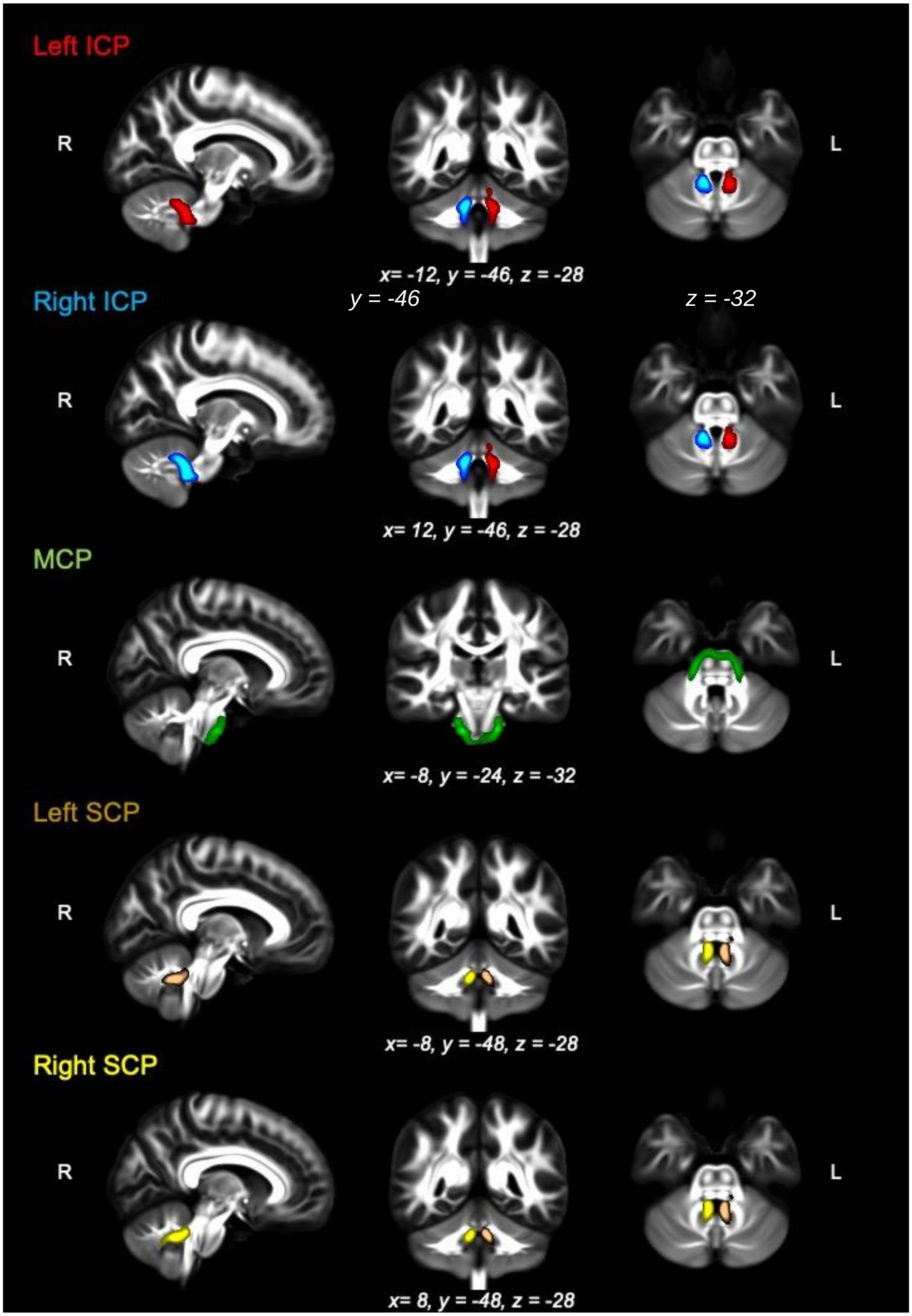

*Supplementary Table (S3)*: Mean tract FA of cerebellar peduncles (CP) in each group

| | TD (N=77) | DLD (N=54) | HSL (N=28) |
|---|---|---|---|
| **Left Inferior CP** | | | |
| Mean (SD) | 0.45 (0.06) | 0.42 (0.06) | 0.44 (0.06) |
| Range (Min - Max) | 0.30 - 0.56 | 0.27 - 0.53 | 0.31 - 0.56 |
| **Right Inferior CP** | | | |
| Mean (SD) | 0.47 (0.04) | 0.45 (0.05) | 0.46 (0.05) |
| Range (Min - Max) | 0.33 - 0.57 | 0.31 - 0.55 | 0.33 - 0.53 |
| **Left Superior CP** | | | |
| Mean (SD) | 0.43 (0.03) | 0.41 (0.03) | 0.44 (0.03) |
| Range (Min - Max) | 0.38 - 0.51 | 0.34 - 0.48 | 0.39 - 0.54 |
| **Right Superior CP** | | | |
| Mean (SD) | 0.42 (0.04) | 0.41 (0.03) | 0.43 (0.03) |
| Range (Min - Max) | 0.34 - 0.51 | 0.34 - 0.49 | 0.37 - 0.48 |
| **Middle CP** | | | |
| Mean (SD) | 0.43 (0.04) | 0.42 (0.04) | 0.42 (0.03) |
| Range (Min - Max) | 0.29 - 0.50 | 0.33 - 0.52 | 0.34 - 0.49 |

*Supplementary Table (S4)*: Model summaries for fractional anisotropy (FA) in the inferior cerebellar peduncles (ICP) when excluding HSL group from the analysis.

| | ICP Model | | | ICP Model including Age, Sex, Motion | | |
|---|---|---|---|---|---|---|
| Predictor | Beta | 95% CI[1] | p-value | Beta | 95% CI[1] | p-value |
| group | | | | | | |
| TD | — | — | | — | — | |
| DLD | -0.09 | -0.17, -0.02 | 0.013 | -0.08 | -0.16, -0.01 | 0.025 |
| hemisphere | | | | | | |
| l | — | — | | — | — | |
| r | 0.09 | 0.03, 0.15 | 0.002 | 0.09 | 0.03, 0.15 | 0.002 |
| whole_brain_FA | 1.6 | -1.8, 5.0 | 0.4 | 1.7 | -1.8, 5.1 | 0.3 |
| group * hemisphere | | | | | | |
| DLD * r | 0.03 | -0.05, 0.12 | 0.4 | 0.03 | -0.05, 0.12 | 0.4 |
| ageInYears | | | | 0.00 | -0.01, 0.02 | 0.6 |
| sex | | | | | | |
| Male | | | | — | — | |
| Female | | | | 0.08 | 0.02, 0.14 | 0.006 |
| relMotion | | | | 0.11 | -0.13, 0.35 | 0.4 |

[1]CI = Confidence Interval
